# TAFFISH: shell-native command-level reproducibility for bioinformatics

**DOI:** 10.1101/2025.09.15.672424

**Authors:** Kaiyuan Han, Ting Wang, Shishi Yuan, Caiyi Ma, Wei Su, Xiaolong Li, Kejun Deng, Hao Lin, Hao Lyu

## Abstract

**Summary:** Bioinformatics analyses often rely on shell commands and small shell scripts whose executable context is difficult to preserve, inspect and reuse. TAFFISH addresses this gap by packaging command-line tool calls and lightweight shell flows as installable, versioned and inspectable executable units. Through a curated public Hub, TAFFISH indexes command interfaces, execution backends, platform constraints, release metadata and smoke-test/validation records. Together, these components provide a command-level reproducibility layer that works directly in ordinary shells and can also be invoked from existing workflow systems.

**Availability and Implementation:** TAFFISH is implemented in Common Lisp and released under Apache-2.0. TAFFISH version 0.10.1, together with the source code, documentation and public Hub resources, is available at https://github.com/taffish/taffish, https://taffish.com, https://taffish.github.io and https://github.com/taffish/. The submitted software release, frozen Hub snapshot, test data, command logs, checksums and reproducibility package are archived at https://doi.org/10.5281/zenodo.21054185. The authors will maintain the software and public resources for at least two years after publication.

**Contact:** **Hao Lyu; Hao Lin; Kejun Deng:**

**Supplementary information:** **Supplementary data are available online.**

## Introduction

Bioinformatics analyses are still routinely assembled from shell commands. On workstations, servers and clusters, researchers invoke tools for quality control, alignment, quantification, sequence processing, phylogenetic inference and reporting, then preserve these steps in scripts, README files, laboratory notes or manuscript methods. This practice is flexible and universal, but important execution semantics often remain implicit. A visible command may depend on a specific software version, container image, language library, system dependency, environment variable, reference dataset or local path, leaving the executable context only partially preserved. TAFFISH starts from the observation that many reproducibility failures arise before a complete workflow exists, at the level of individual command invocations and small shell-based compositions.

Existing workflow systems, package managers, container platforms, descriptor frameworks and environment-capture tools address reproducibility at workflow, software, environment or platform-integration levels (Amstutz et al., 2016; Chirigati et al., 2016; Courtès and Wurmus, 2015; da Veiga Leprevost et al., 2017; Di Tommaso et al., 2017; Dolstra et al., 2004; Glatard et al., 2018; Grüning et al., 2018; Köster and Rahmann, 2012; Kurtzer et al., 2017; The Galaxy Community, 2022). TAFFISH targets a smaller but common layer: individual command-line invocations and lightweight shell flows before they are formalized into complete workflows. It makes the executable command itself an installable and inspectable unit of reuse. Each package binds a shell entry point to its interface, backend and platform constraints, release metadata and validation record, yielding a command callable from ordinary shells, scripts, HPC jobs or workflow processes. A more detailed comparison is provided in Supplementary Table S3.

## System overview

TAFFISH uses a two-level executable package model embedded in the shared shell command pool (Figure 1). At the tool level, a taf-app/tool package bundles an individual command with a .taf interface, version metadata, execution-backend information, platform constraints and basic validation records. For routine analyses, this reduces repeated configuration of container invocations and keeps the command interface and package records together across supported backends. After installation, users obtain directly callable taf-* commands:

~~~
taf update
taf install seqkit
taf-seqkit seqkit stats toy.fa
~~~

**Figure 1.**
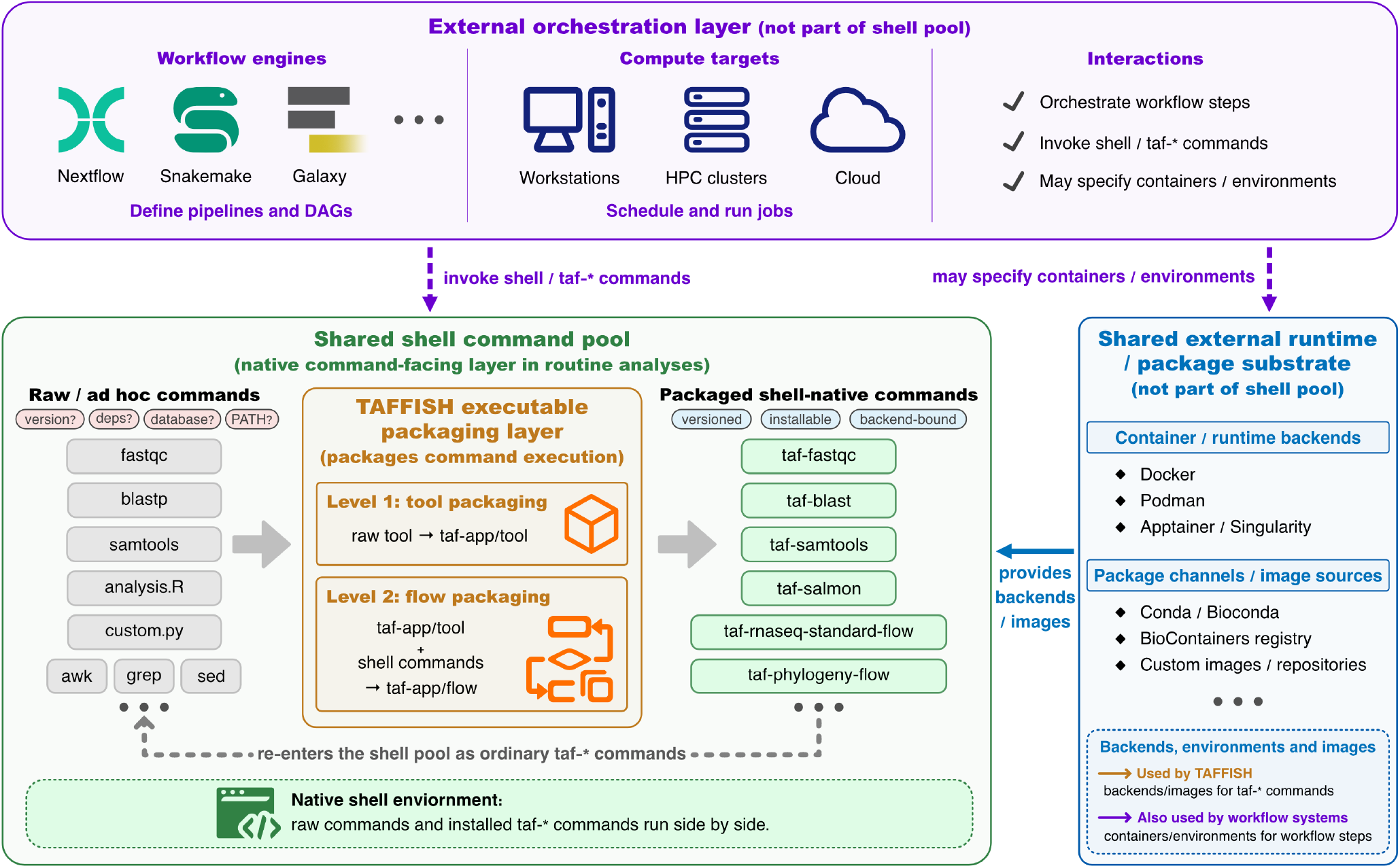
TAFFISH in the bioinformatics command-execution ecosystem. TAFFISH is embedded in the shared shell command pool between ad hoc shell practice, workflow orchestration systems and external runtime/package infrastructure. It packages raw commands and lightweight flows into reproducible taf-app/tool and taf-app/flow objects that re-enter the shell pool as ordinary installable taf-* commands. These commands remain shell-native while using external container/runtime backends and package channels or image sources to make execution portable. *Alt text: Schematic of the TAFFISH ecosystem. Workflow engines and compute targets can invoke shell or taf-* commands. TAFFISH sits inside the shared shell command pool and turns raw tools and lightweight flows into installable shell-native commands. External runtime/package infrastructure provides container/runtime backends, package channels and image sources used by TAFFISH and workflow systems.*

Installed commands can be inspected through their Hub records and the archived JSON/TSV metadata provided with this manuscript. At the package level, taffish.toml records package identity, version/release identifiers, exposed command, backend image, supported platforms and smoke-test metadata, while the .taf source can forward upstream commands or declare options, defaults and required arguments. Users can inspect the selected package with taf-seqkit --version and its generated shell with --compile. Supplementary Note 1b illustrates these interfaces. Unversioned commands select the highest installed version within their installation location. For reproducible scripts, use the explicit version-and-release entry point, such as taf-seqkit-v2.13.0-r2 seqkit stats toy.fa (Supplementary Note 1c).

At the flow level, taf-app/flow packages lightweight shell-native compositions of existing taf-app/tool commands and scripts. A typical use case is the transition from exploratory or one-off shell scripts to shareable reproducible analysis commands without requiring users to first rewrite the analysis in a workflow language.

Such flows are intended to make command execution installable and recordable, not to replace workflow engines that provide DAG scheduling, task caching or cloud/HPC orchestration. Because installed TAFFISH commands remain ordinary shell commands, they can be used in interactive terminals, shell scripts, HPC job scripts or workflow-system processes.

The TAFFISH system consists of the local compiler taffish, the local package manager taf and TAFFISH Hub. The compiler reads .taf sources and generates inspectable shell-level execution logic, while the package manager updates indexes, discovers applications, installs packages and exposes local commands. TAFFISH Hub publishes static records for versions, commands, source/container identities, backend and platform support, smoke-test/validation status and release-integrity evidence such as checksums or signed checksum manifests where available. The archived paper snapshot defines the submitted state, while the live Hub may continue to evolve after submission. TAFFISH 0.10.1 targets POSIX shell and Linux/HPC-oriented bioinformatics use, supports Docker, Podman and Apptainer as execution backends, and provides prebuilt binaries for Linux x86_64 and macOS Apple Silicon. Additional POSIX environments can build from source where the required Lisp and system tools are available. The paper snapshot, generated at 2026-06-29T09:18:17Z, records 200 packages, 305 versions, 200 installable commands and 200 repository records. It contains zero warnings, zero failed records and eight rejected immutable releases documented with replacement versions in Supplementary Data.

### Examples and reproducibility evidence

We use two public examples as evidence for complementary claims. The RNA-seq flow family shows that TAFFISH can package a multi-module, report-generating analysis as installable shell-native flows, with public reference-route and de novo reports and archived run records. The phylogeny-flow example provides a compact clean-install case in which the same cytochrome c analysis route is represented by a public report and by Docker, Podman and Apptainer smoke-test records. Complete commands, versions, inputs, checksums, report links and run manifests are provided in Supplementary Data.

The submitted reproducibility package supports the software availability and reproducibility claims made here. It includes the TAFFISH 0.10.1 release payload, source snapshot, frozen Hub index, supplementary tables, example evidence and backend smoke-test records. Clean-install and runtime-smoke records cover a clean Ubuntu 24.04 x86_64 environment, Linux Docker/Podman/Apptainer execution and representative macOS Docker/Podman tool-level execution. These records are not intended to establish new biological findings or benchmark workflow engines. Package-level smoke tests check selected tool operations, whereas flow tests check a specified multi-tool route and its expected outputs. Passing these tests does not establish general biological correctness or validate untested branches (Supplementary Note 5a).

### Scope and limitations

TAFFISH does not, by itself, guarantee complete reproducibility of scientific conclusions. Results may still depend on input data, references, random seeds, hardware, upstream tools, external databases, container availability and local infrastructure. TAFFISH instead packages command entry points, interfaces, versions, metadata, backend resolution and basic validation records into shell-native executable commands, making routine command execution more portable, inspectable and reusable. The submitted Hub snapshot is maintainer-curated, community package submission is not yet supported, and this work does not claim reproducible builds, SLSA provenance or artifact attestations. Backend portability also does not imply identical permissions, input/output handling or byte-identical results for every analysis (Supplementary Note 5b).

## Data availability

Source code, documentation and public Hub resources are available at https://github.com/taffish/taffish, https://taffish.com,

## Acknowledgements

This work was supported by Center for HPC, University of Electronic Science and Technology of China.

The scientific framing, initial manuscript and Figure 1 were prepared by the authors. OpenAI Codex assisted with revision drafting, language editing, inspection of code and archived records, and local software checks. No new biological analyses were performed during revision. The authors retain responsibility for verifying the technical statements, supporting evidence and final text.

## References

Amstutz,P. et al. (2016) Common Workflow Language, v1.0. Figshare. doi:10.6084/m9.figshare.3115156.v2.

Chirigati,F. et al. (2016) ReproZip: computational reproducibility with ease. In: Proceedings of the 2016 International Conference on Management of Data. ACM, pp. 2085–2088. doi:10.1145/2882903.2899401.

Courtès,L. and Wurmus,R. (2015) Reproducible and user-controlled software environments in HPC with Guix. In: Euro-Par 2015: Parallel Processing Workshops. Lecture Notes in Computer Science. Springer, pp. 579–591. doi:10.1007/978-3-319-27308-2_47.

da Veiga Leprevost,F. et al. (2017) BioContainers: an open-source and community-driven framework for software standardization. Bioinformatics, 33, 2580–2582.

Di Tommaso,P. et al. (2017) Nextflow enables reproducible computational workflows. Nature Biotechnology, 35, 316–319.

Dolstra,E. et al. (2004) Nix: a safe and policy-free system for software deployment. In: Proceedings of the 18th USENIX Large Installation System Administration Conference, pp. 79–92.

Glatard,T. et al. (2018) Boutiques: a flexible framework to integrate command-line applications in computing platforms. GigaScience, 7, giy016.

Grüning,B. et al. (2018) Bioconda: sustainable and comprehensive software distribution for the life sciences. Nature Methods, 15, 475–476.

Köster,J. and Rahmann,S. (2012) Snakemake: a scalable bioinformatics workflow engine. Bioinformatics, 28, 2520–2522.

Kurtzer,G.M. et al. (2017) Singularity: scientific containers for mobility of compute. PLoS ONE, 12, e0177459.

The Galaxy Community. (2022) The Galaxy platform for accessible, reproducible and collaborative biomedical analyses: 2022 update. Nucleic Acids Research, 50, W345–W351.

