## Supplementary Material for "TAFFISH: shell-native command-level reproducibility for bioinformatics"

This supplementary material supports the preprint manuscript by providing implementation details, the frozen software and Hub snapshot, smoke-test/validation records, example reproduction summaries, related-system comparison notes, platform/licensing boundaries and AI-assisted editorial tools disclosure. It is designed as a single human-readable supplementary file for online preprint posting and journal review.

#### Supplementary Material overview

This file contains the supplementary text and summary tables needed to interpret the manuscript-level claims. The complete machine-readable evidence package, including the submitted TAFFISH release payload, Hub snapshot, example manifests, command logs, checksums, release binaries and source archive, is provided separately as a stable external reproduction archive. Paths prefixed with archive: refer to the root of that external archive. The archive DOI is https://doi.org/10.5281/zenodo.21054185.

The material is organized as follows:

| Section | Content | Main purpose |
|---|---|---|
| Supplementary Note 1 | Executable package model and minimal .taf package anatomy | Defines the TAFFISH command-level abstraction used in the main manuscript. |
| Supplementary Notes 2-4 | TAFFISH 0.10.1 release, frozen Hub snapshot and smoke-test/validation records | Documents the submitted software/index state and rejected immutable releases. |
| Supplementary Note 5 | Clean-install and backend smoke-test protocol | Summarizes the installation and runtime evidence supporting software availability. |
| Supplementary Notes 6-7 | RNA-seq flow family and phylogeny-flow examples | Summarizes the two public examples cited in the main manuscript. |
| Supplementary Note 8 | Minimal shell/script/workflow compatibility snippets | Shows that installed taf-* commands remain ordinary shell commands. |
| Supplementary Notes 9-12 | Limitations, AI-assisted editorial tools disclosure, maintenance/release integrity and platform/licensing boundaries | Records scope, disclosure and long-term-use boundaries. |
| Supplementary Tables S1-S5 | Hub app/version lists, validation summary, related-system comparison and evidence manifests | Describes the machine-readable tables mirrored in the external archive. |

**Supplementary Note 1. TAFFISH executable package model**

TAFFISH packages bioinformatics command-line tools and lightweight flows as shell-native executable packages. The model is organized around two app types.

First, a taf-app/tool package represents an individual tool. It combines a command entry point, a .taf interface definition, application metadata, version information, platform/runtime information, container backend information, upstream provenance and basic validation records. After installation, the user obtains a taf-* command that behaves as an ordinary shell command while carrying reproducible execution metadata.

Second, taf-app/flow packages lightweight compositions of existing taf-app/tool packages, shell commands and scripts. Flow apps are intended for installable and reproducible shell-native execution rather than large-scale workflow orchestration. They do not replace workflow systems that provide scheduling, DAG execution, caching or cloud/HPC orchestration.

The core distinction is that TAFFISH packages executable command semantics rather than only software environments or workflow graphs. This allows packaged commands to remain usable in ordinary shells, shell scripts, HPC job scripts and existing workflow systems.

TAFFISH packages are therefore not intended to be simple Docker aliases or a flat wrapper collection. A plain alias can expose a container invocation, but it does not by itself define a versioned package record, a maintained shell entry point, source and container identities, platform constraints, smoke-test status, release-integrity metadata, rejected-release handling or Hub-level indexing. TAFFISH binds these elements into the executable-package record and makes the resulting command installable through taf.

**Supplementary Note 1b. Minimal .taf package anatomy**

A minimal TAFFISH tool package separates application metadata from the executable command interface. For example, the submitted seqkit tool package records package identity, upstream provenance, command name, runtime mode, container image, supported image platforms and smoke-test metadata in taffish.toml, while the .taf source maps user arguments to the containerized command:

```
[package]
name = "seqkit"
kind = "tool"
version = "2.13.0"
release = 2
main = "src/main.taf"

[command]
name = "taf-seqkit"

[container]
image = "ghcr.io/taffish/seqkit:2.13.0-r2"
build_platforms = "linux/amd64,linux/arm64"
```

```
<taf-app:container:ghcr.io/taffish/seqkit:2.13.0-r2>
seqkit ::*ARGV*::
```

This example illustrates the smallest taf-app/tool pattern: a versioned package record, an installable shell command, a resolvable container runtime and a .taf command template. Flow packages use the same executable-package concept but bind multiple installed commands and scripts into a higher-level shell-native command.

Argument forwarding is one interface pattern, not a restriction of the .taf language. An ARGS block can declare long and short options, defaults and required values. The following self-contained example uses shell output to show argument binding without requiring a container:

```
ARGS
<!(--/-i)input>
<(--/-n)label>
sample
RUN
<shell>
printf '%s:%s\n' "::label::" "::input::"
```

Compiling with --input toy.fa and executing the generated shell prints sample:toy.fa; -i toy.fa -n control prints control:toy.fa. Omitting input causes argument binding to fail. These cases were checked locally with the archived 0.10.1 compiler during revision. In contrast, the seqkit example uses taf-app command mode: a leading non-option argument selects an upstream command. In this mode, that command is forwarded rather than checked against a complete TAFFISH declaration of upstream options.

Users can inspect taffish.toml, the .taf source and Hub records for the declared interface, version, source/container identity, platform information and recorded checks. The installed wrapper also exposes package information through --version and generated execution code through --compile, for example taf-seqkit --compile seqkit stats toy.fa. These inspection modes do not establish that every upstream option has been validated.

Detailed .taf DSL syntax, package-writing instructions and backend-specific usage notes are maintained in the TAFFISH online documentation and GitHub repositories rather than reproduced in this Application Note.

**Supplementary Note 1c. Installed versions and reproducible invocation**

TAFFISH 0.10.1 installs an exact version-and-release launcher and an unversioned alias. Installing or removing a version refreshes the alias to the highest remaining installed version for that command in the same installation location. Installing an older version does not make it the default. This is a local selection rule, not a request for the latest online release. If user and system installations coexist, shell PATH order determines which command location is found first.

For scripts that must retain a particular package release, install and call the exact entry point:

```
taf install seqkit 2.13.0-r2
taf-seqkit-v2.13.0-r2 seqkit stats toy.fa
```

The unversioned wrapper reports its selected launcher with taf-seqkit --version. To find the installation path when several versions are present, specify the version, for example taf which seqkit 2.13.0-r2. Pinning the package does not by itself fix the TAFFISH client, backend, input data, parameters, random seeds or external resources.

### Supplementary Note 2. TAFFISH software release used by the manuscript

The manuscript uses TAFFISH version 0.10.1.

Submitted source commit recorded for the manuscript:

```
29f66814f374293223735fa20b7fc74d99aa3ed9
```

Release payload files included in the submitted TAFFISH 0.10.1 archive:

| File | Purpose |
|---|---|
| taffish-darwin-arm64-0.10.1 | TAFFISH compiler binary for macOS Apple Silicon |
| taf-darwin-arm64-0.10.1 | TAFFISH package manager binary for macOS Apple Silicon |
| taffish-linux-amd64-0.10.1 | TAFFISH compiler binary for Linux x86_64 |
| taf-linux-amd64-0.10.1 | TAFFISH package manager binary for Linux x86_64 |
| taffish-mcp-darwin-arm64-0.10.1 | MCP helper binary for macOS Apple Silicon |
| taffish-mcp-linux-amd64-0.10.1 | MCP helper binary for Linux x86_64 |
| SHA256SUMS | release checksum manifest |
| SHA256SUMS.asc | signed checksum manifest |
| TAFFISH-RELEASE-KEY.asc | release public key |

The taffish-mcp helper binary is distributed as part of the TAFFISH 0.10.1 release payload but is not required for the examples or conclusions of this Application Note.

The external reproduction archive prepared for this manuscript contains the submitted source/release evidence, checksum files, Hub snapshot, example records and smoke-test logs. The archive DOI is https://doi.org/10.5281/zenodo.21054185.

### Supplementary Note 3. Frozen paper snapshot of TAFFISH Hub

The manuscript uses the paper snapshot label:

```
paper-01-snapshot-2026-06-29
```

This snapshot records the Hub/index state used by the manuscript. It does not prevent TAFFISH Hub from continuing to grow after submission. Instead, it provides a frozen reference point for the submitted paper.

Snapshot source files:

| Item | URL or file |
| --- | --- |
| Public index URL | <https://raw.githubusercontent.com/taffish/taffish-index/main/index/index.json> |
| Public trust/report URL | <https://raw.githubusercontent.com/taffish/taffish-index/main/index/reports/latest.json> |
| Public Hub pages | <https://taffish.github.io> |
| Archived index copy | archive:evidence/snapshots/2026-06-29/taffish-index.index.json |
| Archived trust/report copy | archive:evidence/snapshots/2026-06-29/taffish-index.latest-report.json |

Snapshot summary:

| Field | Value |
| --- | --- |
| Generated at | 2026-06-29T09:18:17Z |
| Packages | 200 |
| Versions | 305 |
| Commands | 200 |
| Repositories | 200 |
| Warnings | 0 |
| Failed records | 0 |
| Rejected records | 8 |
| Tool versions | 257 |
| Flow versions | 48 |

All package counts, Hub records and validation summaries in the manuscript, Supplementary Data and external reproduction archive refer to this submitted cloud-public snapshot. The example evidence and clean-install/runtime smoke-test records use the same TAFFISH 0.10.1 release and the same RNA-seq and phylogeny-flow demonstrations.

Snapshot checksums:

| File | SHA256 |
| --- | --- |
| taffish-index.index.json | 11d2bb5c254a423609d5d62c024b27d47786f4d5dcd1f1bf104 61ba5c95512da |
| taffish-index.latest-report.json | 293fa2bd328fc2bd1f53a0141d0690e30011dedaf3705d8dba37 bd70bc9b90bb |

**Supplementary Note 4. Hub validation and rejected records**

The snapshot contains zero failed records and zero warnings. Eight immutable releases are rejected and excluded from the main index, each with a replacement version or a documented exclusion reason.

| Repository | Rejected version | Replacement | Short reason |
|---|---|---|---|
| taffish/cooltools | 0.7.1-r1 | 0.7.1-r2 | Docker image build failure |
| taffish/metabat2 | 2.18-r1 | 2.18-r2 | invalid smoke-test metadata |
| taffish/bakta | 1.12.0-r1 | 1.12.0-r2 | smoke-test metadata used an online database-version query |
| taffish/antismash | 8.0.4-r1 | 8.0.4-r2 | Docker image build failure |
| taffish/interproscan | 5.77-108.0-r2 | 5.77-108.0-r3 | incorrect default EBI data archive URL in helper |
| taffish/deepvariant | 1.10.0-r1 | 1.10.0-r2 | invalid smoke-test metadata |
| taffish/muscle | 5.3-r3 | 5.3-r4 | Docker image build failure |
| taffish/fastp | 1.3.3-r1 | 1.3.3-r2 | invalid smoke commands reused a temporary directory |

Tool packages are smoke-tested at the index level with container-command checks, whereas flow packages are validated through package composition plus representative end-to-end runtime smoke tests. Tool-level records in the snapshot contain container images, image digests, smoke-test status and release-integrity metadata. Flow-level records compose existing tool apps and shell commands, and therefore do not have direct container-image smoke-test records at the index level. Representative flow-level evidence is provided separately through the RNA-seq public report-level evidence/checksum records and the phylogeny-flow runtime smoke tests.

**Supplementary Note 5. Clean-install and smoke-test protocol**

The evidence package includes clean-install evidence for Linux x86_64 and runtime smoke-test records using real container backends. Docker, Podman and Apptainer are backend choices; backend availability depends on local infrastructure and permissions. The submitted archive uses TAFFISH 0.10.1 and the paper-01-snapshot-2026-06-29 Hub snapshot, and the clean-install/runtime smoke-test records were generated with the 0.10.1 release binaries.

Clean-install evidence was generated in a clean ubuntu:24.04 amd64 container. After installing ca-certificates, curl and git, TAFFISH 0.10.1 started, updated the public Hub index, resolved phylogeny-flow, installed seqkit 2.13.0-r2, installed phylogeny-flow 0.2.0-r1 and resolved the installed launchers and source records. These logs are provided as:

```
archive:evidence/clean-install/ubuntu2404-amd64-index-smoke.log
archive:evidence/clean-install/ubuntu2404-amd64-install-smoke.log
```

These logs validate binary startup, index update and app installation from a clean Linux user environment.

A separate set of runtime smoke tests was generated on a native Debian x86_64 server using Apptainer, Docker and Podman:

```
archive:evidence/clean-install/linux-apptainer-runtime/linux-
apptainer-environment-probe.log
archive:evidence/clean-install/linux-apptainer-runtime/linux-
apptainer-runtime-smoke.log
archive:evidence/clean-install/linux-apptainer-runtime/linux-
apptainer-cytc-out.sha256
archive:evidence/clean-install/linux-apptainer-
runtime/run.manifest.json
archive:evidence/clean-install/linux-apptainer-
runtime/flow_summary.tsv
archive:evidence/clean-install/linux-apptainer-runtime/versions.tsv
archive:evidence/clean-install/linux-docker-runtime/linux-docker-
runtime-smoke.log
archive:evidence/clean-install/linux-docker-runtime/docker-cytc-
out.sha256
archive:evidence/clean-install/linux-podman-runtime/linux-podman-
runtime-smoke.log
archive:evidence/clean-install/linux-podman-runtime/podman-cytc-
out.sha256
```

The runtime environment was Linux 6.1.0-33-amd64 on x86_64. Each backend smoke initialized an isolated TAFFISH user home, updated the public Hub index, installed seqkit 2.13.0-r2, installed phylogeny-flow 0.2.0-r1 and its dependencies, and ran the 24-sequence cytochrome c example through the MAFFT -> trimAl -> IQ-TREE route. The Apptainer, Docker and Podman logs ended with ## smoke passed and generated output checksums.

Representative macOS Apple Silicon backend smoke-test records were also generated using Docker Desktop and Podman machine:

```
archive:evidence/clean-install/macos-docker-tool-runtime/macos-
docker-tool-smoke.log
archive:evidence/clean-install/macos-docker-tool-runtime/docker-
macos-tool-smoke.sha256
archive:evidence/clean-install/macos-podman-tool-runtime/macos-
podman-tool-smoke.log
archive:evidence/clean-install/macos-podman-tool-runtime/podman-
macos-tool-smoke.sha256
```

These macOS records cover tool-level backend execution from macOS Apple Silicon. The Docker record also includes a representative linux/amd64 DIAMOND command executed through Docker Desktop with DOCKER_DEFAULT_PLATFORM=linux/amd64.

The smoke-test protocol records:

```
taf --version
taffish --version
taf update
taf list
taf install <small-tool-app>
taf-<small-tool-app> --help
taf install phylogeny-flow
taf-phylogeny-flow --help
taf-phylogeny-flow <minimal-test-command>
```

For each run, the evidence package records the operating system, architecture, container backend, backend version, TAFFISH version, Hub snapshot, command logs, exit status, output file manifest and output checksums.

The helper script archive:evidence/clean-install/run-clean-install-smoke.sh prepares the same runtime smoke structure for any Linux environment where Docker, Podman or Apptainer is available.

**Supplementary Note 5a. What the validation records establish**

Hub index validation checks package records and index consistency; it is distinct from executing an analysis. A package-level smoke record reports selected executable and command tests on the recorded backend. For example, seqkit 2.13.0-r2 was checked with Docker for version/help output and small FASTA/FASTQ operations, including conversion, filtering and sequence extraction. A passed record applies to those checks and that environment, not all inputs, options or backends.

The cytochrome c flow tests run MAFFT, trimAl and IQ-TREE together on 24 sequences. They check that the expected input, alignment, tree and report files are nonempty, inspect summary fields and version-specific subcommands, and record output checksums. Tree plotting was disabled. This establishes that the tested route completes and produces the checked output files; it does not validate every optional branch or the biological correctness of arbitrary analyses. The RNA-seq material provides separate public report-level evidence, rather than a new run across all three backends.

**Supplementary Note 5b. Backend portability and output interpretation**

Backend support means that a compatible packaged command can execute through the selected runtime; it does not guarantee identical behavior in every environment. In the archived Linux cytochrome c runs, the checksum manifests agree for 00_inputs/input.fa, 03_results/alignment/aligned.fa, 03_results/alignment/trimmed.fa and 03_results/tree/tree.nwk across Docker, Podman and Apptainer. Other files, including logs, reports and run manifests, are not all byte-identical. This comparison uses the existing archived checksums and is not a new controlled cross-backend experiment.

By default, TAFFISH mounts the host home and working directories and sets the container working directory to the corresponding path. Files outside these locations may need additional mounts, and macOS container virtual machines must be allowed to access the host paths. Permissions remain runtime- and site-dependent. Passing HOME and USER does not set a common numeric UID/GID, and the default Docker/Podman invocation does not add --user. Users should check write access and output ownership in their computing environment.

Simple commands are passed directly to the backend; Docker and Podman use run --rm -i, while Apptainer uses exec. Complex shell blocks can use a bash heredoc, which occupies standard input and should not be assumed to support an independent data stream on stdin. Standard output and error are not normalized across runtimes, and image pulls may emit additional messages. In the archived Docker/Podman flow tests, the images required for the tested route were loaded before analysis. These records support the tested execution paths, but do not establish identical input/output behavior or numerical results in all environments.

**Supplementary Note 6. RNA-seq example**

The first demonstration is a yeast SNF2 bulk RNA-seq flow family.

Public report portal:

<https://taffish.github.io/rnaseq-flows/>

Manuscript-level summary:

| **Field** | **Value** |
|---|---|
| Example type | RNA-seq flow family |
| Public report routes | reference and de novo/no-reference |
| Reference-route samples | 24 |
| Default route | Salmon-first expression route |
| Main modules | reference construction, quantification, differential analysis, enrichment and report generation |
| Optional evidence branches | HISAT2 genome alignment, BAM quality assessment and featureCounts counting |
| Reference-route significant genes reported | 367 |
| Reference-route plots reported | 28 |
| Reference-route HTML reports reported | 52 |

The public portal currently exposes both reference and de novo yeast reports for the same source project. The archive package records the public report entry point, source project, prepared app records, report-flow manifest, key reference-route report metrics and output checksum manifest for this example. It archives report-level evidence rather than the full public HTML output directory.

**Supplementary Note 7. Phylogeny-flow example**

The second demonstration is phylogeny-flow, a compact flow for sequence alignment, optional alignment trimming, phylogenetic tree inference, tree plotting and static report generation. The current public cytochrome c report was generated by phylogeny-flow 0.2.0-r1, matching the submitted paper snapshot. The same compact route is also covered by clean-install Docker, Podman and Apptainer runtime smoke-test records.

Public report:

<https://taffish.github.io/flows/phylogeny-flow/>

Manuscript-level summary:

| Field | Value |
|---|---|
| Example type | phylogenetic inference flow |
| Submitted flow package | phylogeny-flow 0.2.0-r1 |
| Current public report run | phylogeny-flow 0.2.0-r1 example |
| Input | 24 reviewed eukaryotic cytochrome c protein sequences |
| Default route | MAFFT -> trimAl -> IQ-TREE |
| Main outputs | alignment files, trimmed alignment files, Newick tree, tree plots and static HTML report |

Because it is more compact than the RNA-seq example, this case serves as the clean-install reproduction example while still exercising a multi-tool flow.

The archive package records the input FASTA checksum, input source notes, app/version table, runtime commands, environment metadata, output manifest, output checksums, tree output, public report entry point and runtime report output for this example.

**Supplementary Note 8. Minimal shell-compatibility snippets**

Installed TAFFISH commands are ordinary shell commands. Therefore, they can be embedded in shell scripts, HPC job scripts and workflow systems. The following snippets are illustrative shell-compatibility sketches only. They are not counted as independent validation results in this manuscript. The submitted validation evidence is provided by the clean-install records, backend smoke logs, public examples and checksum manifests described in Supplementary Notes 5-7 and Supplementary Table S5.

Shell-script example:

```
taf-samtools samtools --version
taf-seqkit seqkit stats input.fa
```

Nextflow process sketch:

```
process RUN_SEQKIT_STATS {
  input:
  path fasta

  output:
  path "stats.txt"

  script:
  """
  taf-seqkit seqkit stats ${fasta} > stats.txt
  """
}
```

Snakemake rule sketch:

```
rule seqkit_stats:
    input:
        "input.fa"
    output:
        "stats.txt"
    shell:
        "taf-seqkit seqkit stats {input} > {output}"
```

These snippets are intentionally minimal in this Application Note. A future workflow-integration study could evaluate larger Nextflow, Snakemake or HPC scheduler integrations, but that is outside the scope of the present software note.

**Supplementary Note 9. Limitations**

TAFFISH improves reproducible command execution packaging, but it does not guarantee complete reproducibility of scientific conclusions. Results may still depend on input data versions, reference datasets, random seeds, hardware architecture, container image availability, external database updates and upstream tool behavior.

TAFFISH Hub also requires ongoing maintenance. Each app needs accurate metadata, command interfaces, runtime information, basic validation records and version updates. The rejected-record mechanism documents known bad immutable releases but does not eliminate the need for continuous curation.

**Supplementary Note 10. AI-assisted editorial tools disclosure**

The scientific framing, initial manuscript and Figure 1 were prepared by the authors. OpenAI Codex assisted with revision drafting, language editing, inspection of code and archived records, and local software checks. No new biological analyses were performed during revision. The authors retain responsibility for verifying the technical statements, supporting evidence and final text.

The disclosure statement is mirrored in the cover letter and submission metadata.

**Supplementary Note 11. Maintenance, governance and release integrity**

TAFFISH core, taf, TAFFISH Hub records and public documentation are maintained through the taffish GitHub organization. The manuscript uses TAFFISH 0.10.1 and the frozen paper snapshot paper-01-snapshot-2026-06-29; the live Hub may continue to grow after submission, while the submitted manuscript refers to the archived snapshot.

In the submitted version, TAFFISH Hub is a maintainer-curated public index. New package records and version updates are prepared, checked and released by the TAFFISH maintainers. Curation currently checks package identity, upstream source or release information, command interface, container/backend binding, platform declarations, smoke-test metadata, license notes where available and release-integrity metadata. In this manuscript, trust metadata is used narrowly to mean package-level source/container identity, checksums or signed checksum manifests where available, smoke-test status, package-validation status and rejected-release records; it does not imply supply-chain attestations, reproducible builds, SLSA compliance or complete provenance guarantees. Records are included in the main index only after compilation/index validation succeeds and required smoke-test/validation and release-integrity metadata are present; known bad immutable releases are rejected rather than silently replaced. A broader community submission workflow and open platform governance are planned directions, but direct community package submission is not supported in the submitted release.

TAFFISH Hub uses immutable repository/tag/release identities for app versions. Rejected immutable releases are excluded from the main index and replacement versions are documented in the Hub validation report. The submitted release payload includes SHA256SUMS, SHA256SUMS.asc and TAFFISH-RELEASE-KEY.asc. Manual checksum and signature verification is supported for users requiring higher assurance. This submission does not claim reproducible builds, SLSA provenance or artifact attestations.

Issues and maintenance requests are tracked through the public GitHub repositories. The maintainers intend to keep the submitted software release, archived reproducibility package, paper snapshot, public documentation and example entry points available for at least two years after publication.

**Supplementary Note 12. Platform and licensing boundaries**

TAFFISH 0.10.1 provides prebuilt client binaries for Linux x86_64 and macOS Apple Silicon. Packaged tool execution depends on a local container backend and the platform support of each tool image. In the submitted Hub snapshot, 257 tool-version records list linux/amd64 support, 181 tool-version records list linux/arm64 support, and 48 flow-version records have no direct container platform because they compose existing tool apps and shell commands.

TAFFISH package metadata and wrappers are released under Apache-2.0 unless otherwise stated. Upstream tools, databases, reference data and container contents retain their original licenses and redistribution terms.

**Supplementary Tables**

The tables below summarize the machine-readable supplementary tables generated for this manuscript. For journal submission, the primary supplementary material is this single Supplementary Data file. The full TSV files are available under supplementary-tables/, duplicated under archive:supplementary-tables/ in the external reproduction archive and may also be supplied as machine-readable supplementary files if requested by the journal or reviewers.

**Supplementary Table S1. Full app list in the submitted Hub snapshot**
Provided as:

```
supplementary-tables/table-s1-hub-app-list.tsv
```

This table was generated from:

```
archive:evidence/snapshots/2026-06-29/taffish-index.index.json
```

It contains one row per latest app package in the paper snapshot, plus a header row. The generated table has 200 app records.

The TSV header records package identity, latest version, kind, command, repository/source reference, container image and digest, platforms, the smoke_status and trust_status columns for package-level smoke and release-integrity checks, upstream information, license, categories and summary.

**Supplementary Table S1b. Full version list in the submitted Hub snapshot**
Provided as:

```
supplementary-tables/table-s1b-hub-version-list.tsv
```

This table contains one row per version record in the paper snapshot, plus a header row. The generated table has 305 version records.

The TSV header records package identity, version identifier, latest-version flag, command, repository/source reference, container image and digest, platforms, smoke_status and trust_status columns with timestamps for package-level smoke and release-integrity checks, upstream information, license, categories and summary.

**Supplementary Table S2. Hub validation summary**

Provided as:

```
supplementary-tables/table-s2-hub-validation-summary.tsv
```

Summary:

| Category | Count |
|---|---|
| Packages | 200 |
| Versions | 305 |
| Tool versions | 257 |
| Flow versions | 48 |
| Commands | 200 |
| Repositories | 200 |
| Tool versions with passed smoke-test records | 257 |
| Tool versions with container digests | 257 |
| Failed records | 0 |
| Warning records | 0 |
| Rejected records | 8 |

**Supplementary Table S3. Related-system comparison**

Provided as:

```
supplementary-tables/table-s3-related-system-comparison.tsv
```

This table compares abstraction layer, representative systems, primary object, primary interface, relationship to shell and TAFFISH distinction.

**Supplementary Table S4. Example command/version/checksum manifest**

Provided as:

```
supplementary-tables/table-s4-example-evidence-manifest.tsv
```

This table records the current evidence items for the RNA-seq and phylogeny-flow examples, including public report URLs, app versions, archived manifest sources, runtime smoke-test records and generated checksum files.

The TSV header records the example name, evidence item, value, source file or URL, status and note.

**Supplementary Table S5. Backend and clean-install smoke summary**
Provided as:

```
supplementary-tables/table-s5-backend-smoke-summary.tsv
```

This table summarizes clean-install, app-install and representative backend runtime smoke evidence, including operating system, architecture, backend, operation, result and evidence files.
